# Increased processing of SINE B2 non coding RNAs unveils a novel type of transcriptome de-regulation underlying amyloid beta neuro-pathology

**DOI:** 10.1101/2020.07.20.212886

**Authors:** Yubo Cheng, Babita Gollen, Luke Saville, Christopher Isaac, Jogender Mehla, Majid Mohajerani, Athanasios Zovoilis

**Author notes:** Equal contribution (Yubo Cheng, Babita Gollen, Luke Saville, Christopher Isaac).

## Abstract

More than 97% of the mammalian genome is non-protein coding, and repetitive elements account for more than 50% of noncoding space. However, the functional importance of many non-coding RNAs generated by these elements and their connection with pathologic processes remains elusive. We have previously shown that B2 RNAs, a class of non-coding RNAs that belong to the B2 family of SINE repeats, mediate the transcriptional activation of stress response genes (SRGs) upon application of a stimulus. Notably, B2 RNAs bind RNA Polymerase II (RNA Pol II) and suppress SRG transcription during pro-stimulation state. Upon application of a stimulus, B2 RNAs are processed into fragments and degraded, which in turn releases RNA Pol II from suppression and upregulates SRGs. Here, we demonstrate a novel role for B2 RNAs in transcriptome response to amyloid beta toxicity and pathology in the mouse hippocampus. In healthy hippocampi, activation of SRGs is followed by a transient upregulation of pro-apoptotic factors, such as p53 and miRNA-34c, which target SRGs creating a negative feedback loop that facilitates transition to the pro-stimulation state. Using an integrative RNA genomics approach, we show that in mouse hippocampi of an amyloid precursor protein knock-in mouse model and in an *in vitro* cell culture model of amyloid beta toxicity, this regulatory loop is dysfunctional due to increased levels of B2 RNA processing, constitutively elevated SRG expression and high p53 levels. Evidence indicates that Hsf1, a master regulator of stress response, mediates B2 RNA processing in cells, and is upregulated during amyloid toxicity accelerating the processing of SINE RNAs and SRG hyper-activation. Our study reveals that in mouse, SINE RNAs constitute a novel pathway deregulated in amyloid beta pathology, with potential implications for similar cases in the human brain, such as Alzheimer’s disease (AD). This data attributes a role to SINE RNA processing in a pathological process as well as a new function to Hsf1 that is independent of its transcription factor activity.

## INTRODUCTION

The number of patients with Alzheimer’s disease (AD) is expected to skyrocket in upcoming years (Cornutiu, 2015). The exact molecular mechanisms underlying AD are not fully understood, a fact that is underlined by the inauspicious results of recent clinical trials for potential therapeutic agents (Cummings, Lee, Ritter, Sabbagh, & Zhong, 2019; Hane et al., 2017). Amyloid pathology, and particularly, amyloid beta peptides and their aggregated forms have been connected with AD pathogenesis (Bloom, 2014) as well as with neurotoxicity in mouse models of amyloid pathology (Ittner et al., 2010). Nevertheless, the transcriptome changes involved in cell stress response to amyloid toxicity in brains with extensive amyloid beta pathology are still not entirely clear. The hippocampus is a primary target of amyloid pathology in humans. In healthy hippocampi, among the genes that have been implicated in transcriptome-environment interactions are stress response genes (SRGs) (Gallo, Katche, Morici, Medina, & Weisstaub, 2018). These genes have been initially described in other biological contexts, such as thermal and oxidative stress, as pro-survival genes activated early after the application of a stress stimulus, such as heat shock, that help the cell overcome the stress condition (Mahat, Salamanca, Duarte, Danko, & Lis, 2016). However, in addition to the cellular response to stress, many SRGs were shown to have a central role in the function of the mouse hippocampus, by mediating cell signalling and genome-environment interactions. In particular, we and others have shown that in the healthy hippocampus during neural response to environmental stimuli, SRGs, such as those of the MAPK pathway, are transiently activated (Peleg et al., 2010). Activation of SRGs is followed by a transient upregulation of the pro-apoptotic factor p53 and, subsequently, of a pro-apoptotic miRNA, miR-34c, which are transiently induced by and as a response to the activation of pro-survival SRGs (Zovoilis et al., 2011). p53 activates the expression of genes engaged in promoting cell death in response to multiple forms of cellular stress including miRNA-34c (Yamakuchi & Lowenstein, 2009). This miRNA acts transiently as a guard and fine tuner of the expression of many SRGs by targeting them, thus creating a negative feedback regulatory loop that keeps SRG expression in healthy cells under strict control. This facilitates the return to the pro-stimulation state and normalization of p53 levels in approximately 3 hours after the application of the stimulus (Zovoilis et al., 2011). In contrast, hippocampi of mouse models of amyloid pathology and post-mortem brains of human patients of AD are characterized by abnormally high miR-34c levels that subsequently can lead to prolonged high p53 levels and neural death (Zovoilis et al., 2011). Given that many SRGs are upstream regulators of p53-miR-34c activation (Gao et al., 2010), high p53 and miR-34c levels in amyloid pathology implied a possible transcriptome deregulation of the pathways that involve SRGs but whether such a deregulation exists, and which is the mechanism underlying this, it remained unknown. Interestingly, in a recent publication, we showed that expression of a number of SRGs is regulated by a class of non-protein coding (non-coding) RNAs called B2 SINE RNAs (Zovoilis, Cifuentes-Rojas, Chu, Hernandez, & Lee, 2016), raising the possibility that these non-coding RNAs may be a missing link in the pathways connecting amyloid beta toxicity with transcriptome changes in mouse hippocampus during amyloid pathology.

SINE non-coding RNAs are transcribed by repetitive Small Interspersed Nuclear Elements (SINEs), with the subclass B2 repetitive elements being one of the most frequent in mouse (Kramerov & Vassetzky, 2011). SINE B2 elements, which are retrotransposons present in millions of copies, are part of the non-protein coding genome, and over the long haul they have been regarded as genomic parasites and “junk DNA” with no function (Karijolich, Zhao, Alla, & Glaunsinger, 2017). However, SINE B2 elements can be transcribed by RNA Polymerase III into SINE B2 RNAs (Kramerov & Vassetzky, 2011) and a number of recent studies have revealed a key role for SINE B2 RNAs in cellular response to stress. In particular, studies from the J.Kugel and J. Goodrich labs have shown that during response to cellular stress, levels of SINE B2 RNAs increase and suppress the transcription of a number of housekeeping genes through binding of RNA Polymerase II (RNA Pol II), potentially facilitating the redirection of cell resources to pro-survival pathways (Yakovchuk, Goodrich, & Kugel, 2009) (Yakovchuk et al., 2009). In addition, we have recently shown that SINE B2 RNAs mediate cellular response to stress through the regulation of pro-survival stress response genes by acting as transcriptional switches. In particular, in the pro-cellular stress state SINE B2 RNAs bind RNA Pol II at several SRGs and suppress their transcription. In this way, stalled or delayed RNA Pol II remains poised for a fast activation and ramp up of transcription when needed. Upon application of a stress stimulus, SINE B2 RNAs, which have a self-cleavage activity that is accelerated by their interaction with some proteins (Hernandez et al., 2020), are processed into unstable fragments that lack the ability to bind and suppress RNA Pol II. This event releases the delayed or stalled RNA Pol II and enables fast transcriptional activation of stress response genes (Zovoilis et al., 2016).

The above findings have revealed a novel role in cellular function for processing of SINE B2 RNAs (hereafter referred simply as B2 RNAs) through the regulation of SRGs. However, an association of the processing and destabilization of B2 RNAs with any pathological cellular process remains unknown. Here, given the importance of SRGs in hippocampal neuronal function, we examine whether this newly described B2 RNA-SRG regulatory mechanism is linked with pathological processes, focusing on transcriptome response to amyloid beta toxicity and pathology. To this end, we investigate whether B2 RNA regulated SRGs are indeed deregulated during amyloid pathology, which would imply a role for B2 RNAs in this condition and we examine whether amyloid toxicity is connected with changes in B2 RNA processing. Subsequently, we investigate the further upstream molecular mechanisms underlying any potential deregulation of B2 RNA processing in response to amyloid toxicity in hippocampal neural cells.

## RESULTS

### B2 RNA regulated SRGs are enriched in neural functions

In a previous study, we have identified genomic locations that are subject to regulation by SINE B2 non-coding RNAs through binding and suppression of transcription by the RNA Pol II at the pre-stimulus state. Upon induction of cellular stress through the application of a stimulus, SRGs in these locations become activated through B2 processing and release of RNA Pol II suppression (Fig.1A) (Zovoilis et al., 2016). A list of B2 RNA regulated SRGs in these locations is available in Suppl. Table 1.

**Figure 1.**
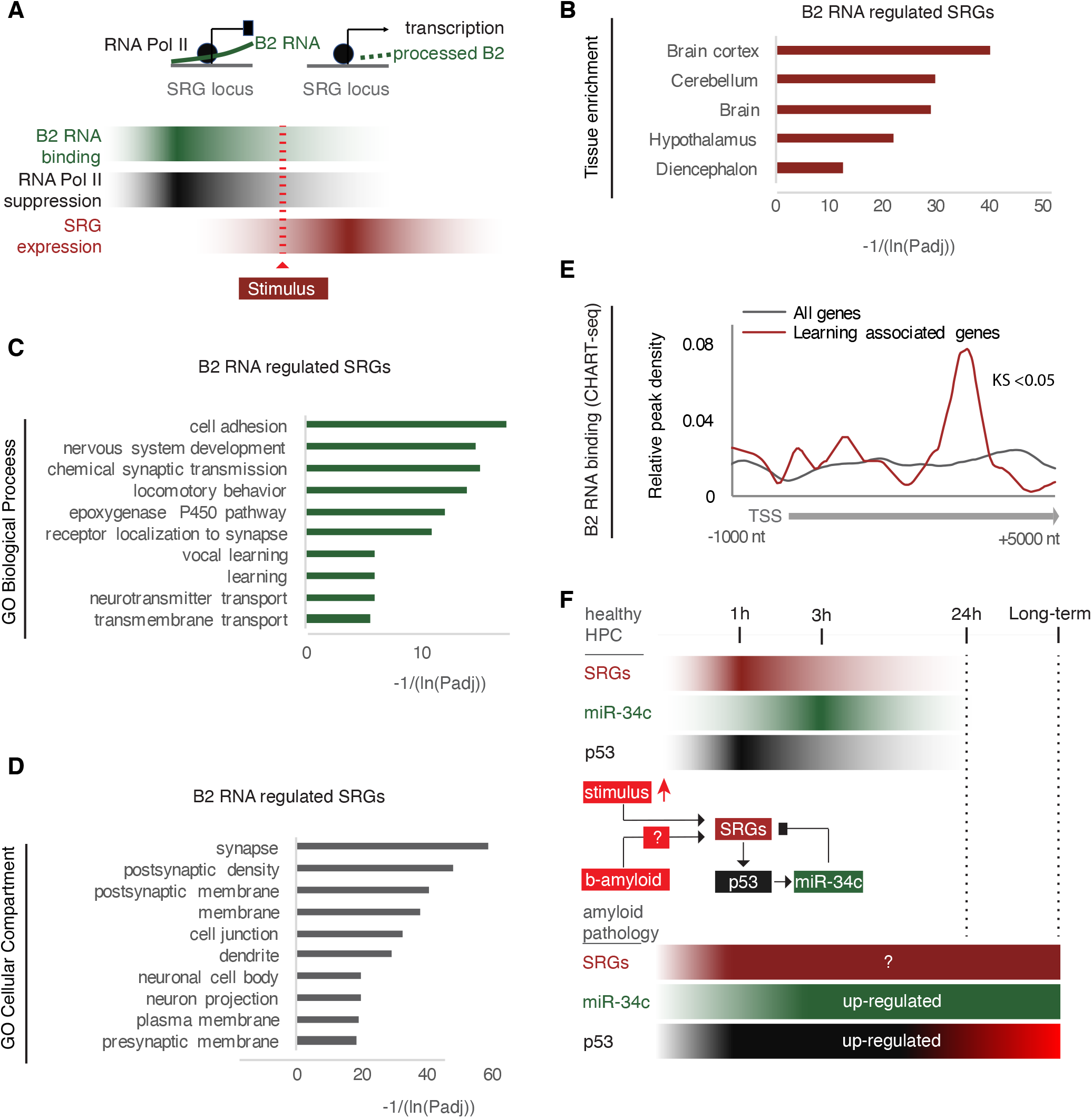
B2 RNA regulated SRGs are enriched in neural functions. **(A)** Regulation mode of SRGs by B2 RNA processing based on previous works (Ponicsan, Kugel, & Goodrich, 2010; Yakovchuk et al., 2009; Zovoilis et al., 2016). Color intensity represents higher B2 RNA binding (green), Pol II suppression (black) or SRG expression (red). **(B)** Tissue enrichment analysis of B2 RNA regulated SRGs listed in Suppl. Table1 based on the DAVID functional annotation platform (Huang da et al., 2009a, 2009b). The adjusted p-values of top ranking terms are plotted as a function with higher scores representing lower adjusted p-values (higher statistical significance) ranging from padj 1.92E-04 for “Diencephalon” to padj 8.91E-13 for “Brain Cortex”. **(C)** Gene ontology (GO) analysis (DAVID) of B2 RNA regulated genes on the basis of cellular compartments. The adjusted p-values of top ranking terms are plotted as a function with higher scores representing lower adjusted p-values ranging from 2.23E-06 for “Presynaptic membrane” to 1.88E-18 for “Synapse”. **(D)** GO analysis (DAVID) of B2 RNA regulated genes on the basis of biological processes. As above, adjusted p-values of top ranking terms are plotted as a function, with higher scores representing lower adjusted p-values ranging from 0.02 for “Transmembrane transport” to 5.44E-06 for “Cell adhesion”. **(E)** Metagene analysis of distribution of genomic B2 RNA binding sites across the start of genes from (Zovoilis et al., 2016), comparing learning associated genes from (Peleg et al., 2010) with all genes (Kolmogorov Smirnov test, KS< 0.05). TSS represents the Transcription Start Site of these genes. **(F)** Regulatory loop of SRGs-p53-miR-34c in mouse hippocampus based on (Zovoilis et al., 2011). Role of amyloid beta in affecting this loop remained unknown after these works (noted with a question mark).

We questioned whether there are known cellular functions connected with B2 RNA regulated SRGs. As shown in Fig. 1B, after performing a tissue enrichment analysis to identify tissue terms that are over-represented in the list of our SRGs, we found a significant enrichment of neural tissue terms compared to other tissues in our list. Similarly, during Gene Ontology term enrichment analysis, cellular compartments closely related to neural functions top the list of enriched terms in these genes, including terms such as synapse, postsynaptic density and membrane, dendrites, neural projections and pre-synaptic membrane (Fig.1C). Most importantly, after performing the same analysis for Biological Processes GO terms, B2 RNA regulated SRGs were found to be enriched considerably in neural function related terms. GO terms, such as learning, nervous system development, synaptic transmission, synapse receptor localization and neurotransmitter transport were among the first 10 entries with the highest adjusted P value scores enriched in B2 RNA regulated SRGs (Fig.1D).

Among the biological processes potentially affected by B2 RNAs that were identified above was learning. We and others have already shown that learning processes in the mouse hippocampus are connected with the transient activation of a number of learning associated genes and pathways, including many known SRGs, (Peleg et al., 2010). For this reason, we examined whether learning associated genes are among the binding targets of B2 RNAs. To this end, we compared the distribution of B2 RNA binding sites in the genome between learning associated genes and all genes. Indeed, as shown in Fig.1E, B2 RNA binding sites were found to be enriched in learning associated genes compared to other genes.

This data suggests that B2 RNA regulated SRGs could affect a wide spectrum of neural functions including learning. Learning is among the biological processes heavily impaired in AD. Given the role of hippocampus as a primary target of amyloid pathology in AD, we decided to focus further on the potential role of B2 RNA-SRGs regulation in this pathological process.

### B2 RNA regulated SRGs get hyper-activated during progression of amyloid pathology

We have previously shown that in hippocampus the pro-apoptotic miRNA miR-34c, which is induced among others by p53, targets in a negative feedback loop manner many SRGs and helps to restore SRG and p53 expression levels after application of a stimulus (Fig.1F) (Zovoilis et al., 2011). In both mouse models of amyloid beta pathology and AD patient brains, we found persistently high levels of miR-34c. High miR-34c levels in hippocampi with amyloid pathology are indicative of a transcriptome-wide deregulation of p53 and SRG levels, as many of these genes are either direct or indirect upstream regulators of miR-34c (Fig.1F) (Rokavec, Li, Jiang, & Hermeking, 2014). Thus, we questioned whether transcriptome changes in response to amyloid pathology involve SRGs regulated by B2 RNAs.

To test whether B2 RNA regulated SRGs are indeed de-regulated in amyloid pathology, we employed a transgenic mouse model of amyloid pathology, APP^NL-G-F^,(Saito et al., 2014) and the respective wild type control (C57BL/6J). The same animal cohorts that were previously characterized through a battery of immunohistochemistry (IHC) and behavioral tests (Mehla et al., 2019) were used to isolate whole hippocampi for the transcriptome analysis conducted in the current study. Fig.2A depicts our experimental design while Fig.2B and Suppl. Fig.1 depict the amyloid plaque deposition in the brains and the behavioral tests, respectively, in these mouse cohorts.

**Figure 2.**
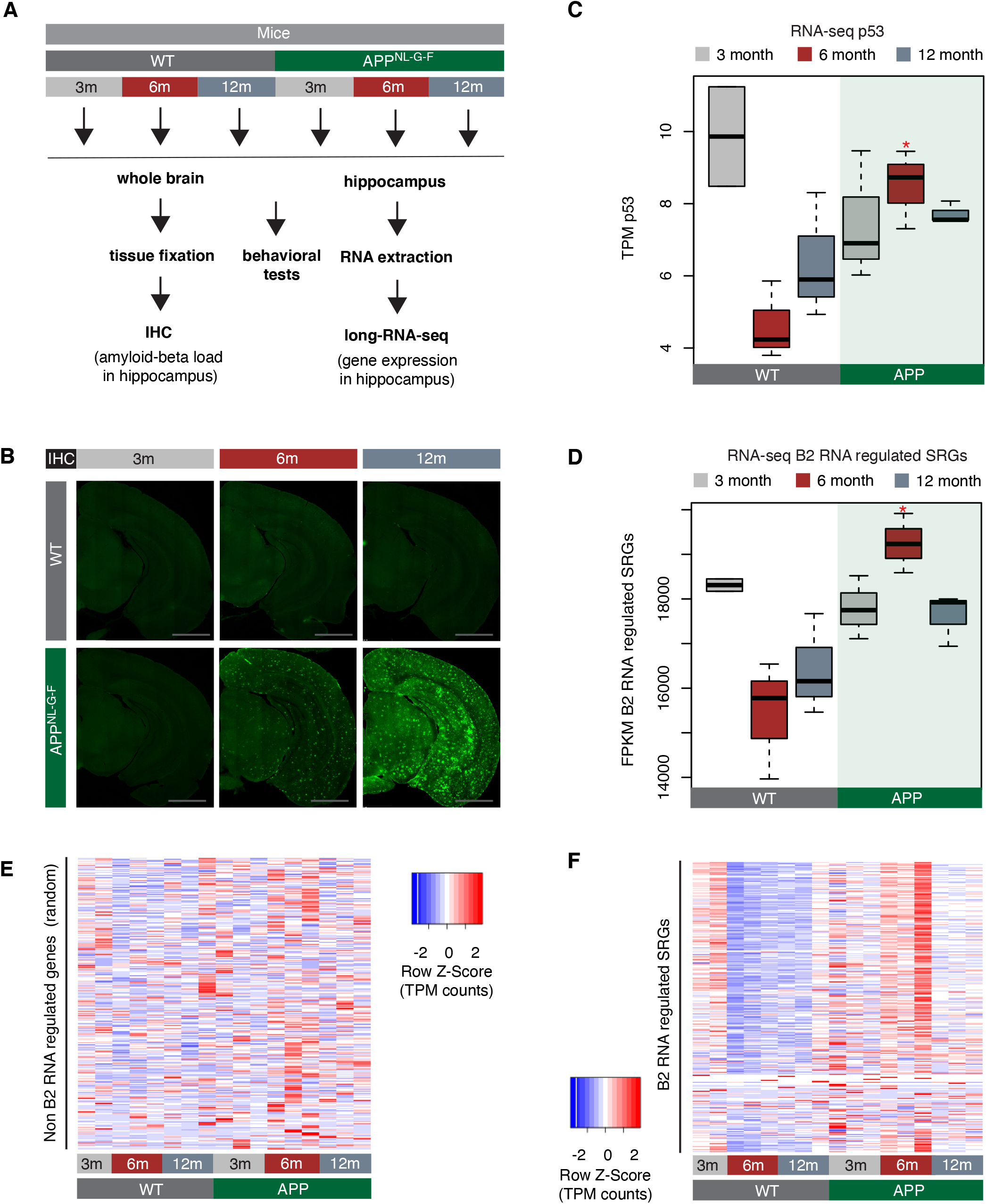
B2 RNA regulated SRGs are hyper-activated in amyloid pathology. **(A)** Experimental design for study of B2 RNA regulated SRGs in the hippocampus of the amyloid pathology mouse model (APP) and the respective wild type (WT) control. **(B)** Immunohistochemistry for identifying amyloid-beta load in the brain sections of mice from the same cohort as in (A) (image credit: Mehla and colleagues, adapted from previously published figure (Mehla et al., 2019)). Higher fluorescence intensity corresponds to higher amyloid load in APP mice from 6m old onwards. **(C)** Expression levels of the pro-apoptotic gene p53 (official symbol TrP53) as defined by long-RNA-seq. p53 transcripts per million reads (TPM) from the APP mice compared to WT, grouped by age. Boxplot depicts distribution of expression levels of p53 gene among different age groups of mice (WT: wild type, APP: mice with amyloid pathology). Black line denotes median. p < 0.05 between 6m old APP and 6m old WT (depicted as asterisk, t-test, n=3/group). **(D)** Expression levels of B2 RNA regulated SRGs as defined by long-RNA-seq. Boxplot depicts distribution of expression levels (in FPKM / Fragments per Kilobase per Million) of B2 RNA regulated genes among different age groups of wild type and APP mice. p < 0.05 between 6m old APP and 6m old WT (depicted as asterisk, t-test, n=3/group). **(E)** Gene expression levels (defined by long-RNA-seq) of a random set of genes not associated with B2 RNAs show weak or no association with amyloid pathology status in the hippocampus of APP and WT mice of different ages. Heatmap depicts gene expression with rows representing 1426 random genes (non B2 RNA regulated) and columns representing the different mouse samples. Expression values are normalized per row and correspond to TPM values. **(F)** Gene expression levels of B2 RNA regulated SRGs (Suppl. Table1) show strong association with amyloid pathology status in the hippocampus of APP and WT mice of different ages. Heatmap depicts gene expression with rows representing B2 RNA regulated genes and columns representing mouse samples. Expression values are normalized per row and correspond to TPM values. Red color represents higher expression.

We have focused on three different mouse ages that correspond to different phases of amyloid beta pathology and represent the: i) pre-symptomatic stage with undetectable (very low) amyloid plaque load, (3 months - 3m, Fig.2B left panels), ii) stage of symptom manifestation (6 months - 6m) (Suppl. Fig.1) that coincides with the active neurodegeneration phase and appearance of amyloid plaques (Fig.2B, middle-panels), and iii) terminal stage of the pathology (12 months - 12m, Fig. 2B right panels) when mice have already acquired the extensive brain atrophy due to neural cell death. Whole hippocampi from mice of these three groups were isolated and the extracted RNA was subjected to next-generation sequencing. We performed directional RNA sequencing for these samples and subsequently quantified gene expression levels.

In accordance with increased cell death during the active neurodegeneration phase at 6 months, levels of p53 were elevated in APP mice of this age compared to controls of the same age (t-test, *p<0.05*, n=3/group) (Fig.2C). Moreover, consistent with our hypothesis of a transcriptome-wide deregulation, levels of B2 RNA regulated SRGs were found to be strongly upregulated during the active phase of neurodegeneration at 6 months (Fig.2D-F). In particular, in healthy hippocampi (WT control animals), levels of these genes are normally downregulated in 6m and 12m old mice compared to 3m old mice *(p=0.02 and p=0.04, respectively)* (Fig.1A and 1F). This is in accordance with the higher level of neural synaptic activity and plasticity that younger mice have (Lilja et al., 2013). In contrast, levels of B2 RNA regulated SRGs in APP mice remain abnormally high in 6m old APP mice compared to both the 3m old APP mice and the 6m old control mice (*p<0.05* in both cases) (Fig.2D and 2F). This pattern in B2 RNA regulated SRG expression was not observed when a random set of genes was tested (Fig.2E).

These findings show a transcriptome wide hyper-activation of B2 RNA regulated SRGs during the active phase of amyloid pathology.

### B2 RNA processing rate increases upon progression of *in vivo* amyloid pathology

Given the role of B2 RNAs in the regulation of SRGs, we hypothesized that the observed hyper-activation of B2 RNA regulated SRGs in the hippocampus of 6m old APP mice may reflect a similar upstream deregulation at the level of the B2 RNAs.

To test this, we employed a customized version of RNA sequencing and analysis used in our previous study (Zovoilis et al., 2016) that allows for enrichment and sequencing of the short SINE RNA fragments (<100nt) produced by B2 RNA processing (short-RNA-seq). In contrast to standard long-RNA-seq protocols that include RNA fragmentation and, thus, may introduce bias, this approach circumvents this problem. Moreover, the long-RNA-seq protocols exclude short RNA fragments of <100nt making the identification of B2 short fragments challenging. After short-RNA-seq (Fig.3A), mapping of the 5’ends of the sequenced fragments across the B2 loci enables the determination of processing points at B2 RNA (depicted as “X” in Fig.3A), including those at the critical RNA Pol II binding region (depicted as a rectangle in Fig.3A). Fig.3B depicts an example of these fragments from one of the samples. As shown in Fig.3A, B2 RNA is extensively processed in hippocampi with amyloid pathology. In particular, this data revealed an increased number of B2 RNA fragments in 6m old mice compared to controls (Fig.3A), a difference not observed when comparing full length B2 RNA levels between these groups of mice using long-RNA-seq (Fig.3C).

**Figure 3.**
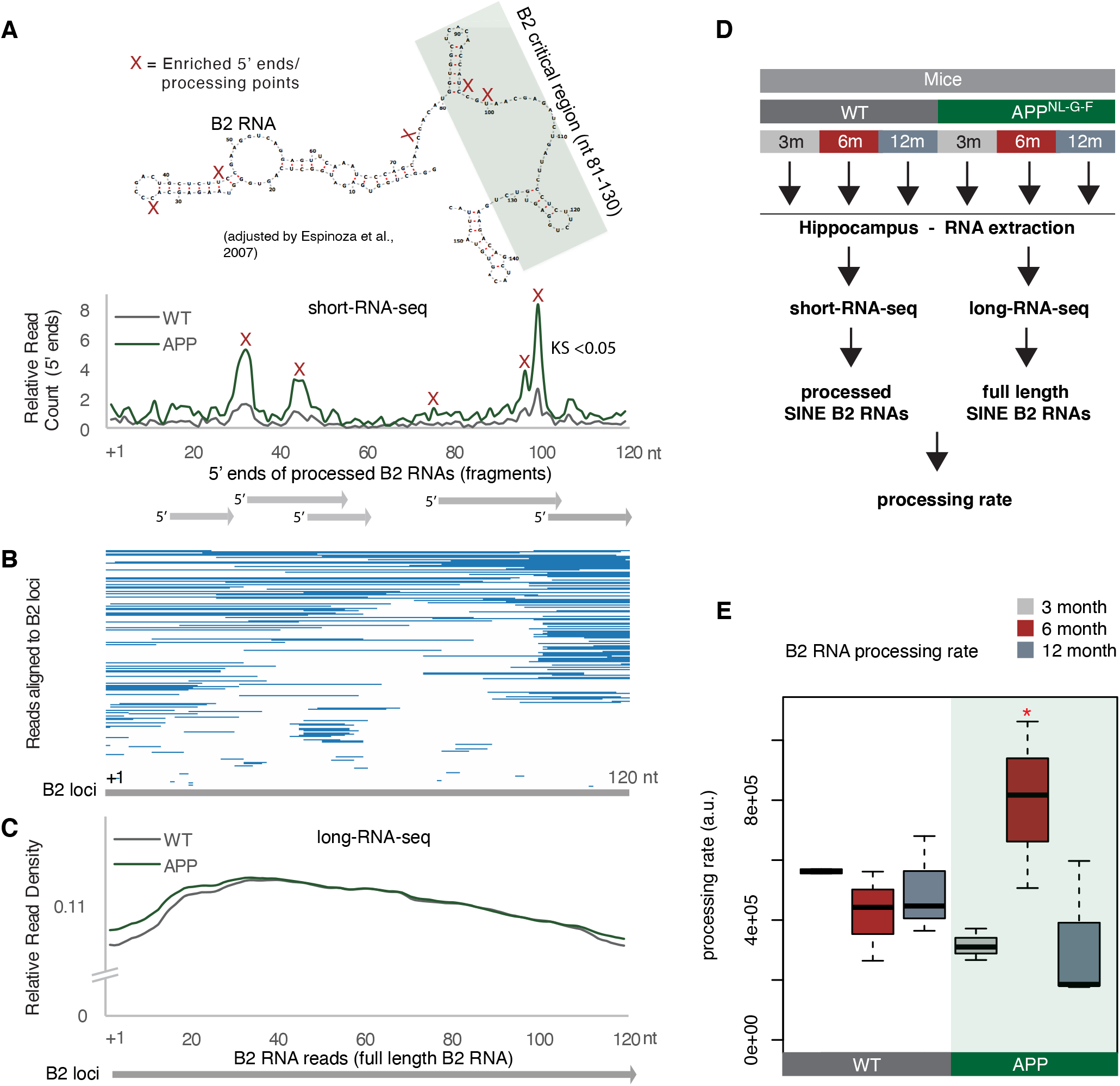
B2 RNA processing rate is increased in 6m old APP mice. **(A)** Plotting of the position of the first base (5’ end) of B2 RNA fragments across the B2 loci to depict increased levels of B2 RNA fragments in 6m old APP mice. **Upper panel**: Secondary structure and processing points of B2 RNA. Secondary structure of B2 RNA adapted from Espinosa and colleagues (Espinoza, Goodrich, & Kugel, 2007). As in our previous study (Zovoilis et al., 2016), we depict the B2 RNA processing points based on short RNA-seq data and mapping of the 5’ends of B2 RNA fragments. X marks which cleavage sites of B2 RNA in the upper panel correspond to enriched processing points (the peaks of 5’end fragments distribution) at the lower panel. The green rectangle depicts the critical region that binds and suppresses RNA Pol II (Espinoza et al., 2007; Ponicsan et al., 2010; Ponicsan, Kugel, & Goodrich, 2015; Yakovchuk et al., 2009) that may be affected by such processing points. **Lower panel:** Distribution of the 5’ends of B2 RNA fragments across the B2 RNA loci based on mapped short RNA-seq from the hippocampi of 6m old APP and WT mice. The x axis represents a metagene combining all B2 RNA loci aligned at the start site of their consensus sequence (position +1) and the y axis shows the relative 5’ end count for B2 RNA fragments aligning to any position downstream of position +1. Figure depicts the difference between B2 RNA fragment distribution of 6m old APP mice (higher peaks) vs 6m old WT mice (lower peaks) (Kolmogorov Smirnov test, KS<0.05). **(B)** Alignment of short-RNA seq reads mapped against the multiple B2 loci. Only B2 loci with at least one read are depicted. Short RNA-seq reads from one of the 6m old APP mice, whose 5’ends are plotted at (A) lower panel are used here as an example of the length and position of fragments across B2 loci. **(C)** Metagene analysis of long-RNA-seq reads of mice across B2 RNA loci that is enriched in full length B2 RNAs but not in B2 RNA fragments. Comparative analysis shows low difference in relative read density of full length B2 RNAs between WT and 6m old APP. X axis as in A. Y axis corresponds to read coverage across the B2 loci. **(D)** Experimental design for estimation of B2 RNA processing rate based on short and long-RNA seq data for all APP mice and controls. **(E)** Boxplot depicts distribution of SINE B2 RNA processing rate among different age groups of mice between wild type and APP. p < 0.05 between 6m old APP and 6m old WT (depicted as asterisk, n=3/group, t-test).

This data suggests that the processing rate of B2 RNAs may be higher in APP mice of this age. To estimate this rate, we used the short RNA-seq data in combination with standard long-RNA-seq (Fig.3D). As in our previous study, for normalization of short RNA 5’end values we used a class of short RNAs that is not affected by B2 RNAs, the RNA Pol III-transcribed tRNAs. Moreover, the absolute numbers of fragmented B2 RNA may vary according to the underlying expression of full length B2 RNA transcripts. Therefore, in order to factor in any differences in the amount of fragments due to basal expression levels of the full length B2 RNA, B2 RNA fragment values from short RNA-seq were normalized to the levels of the full length B2 RNAs calculated by the directional long-RNA-seq. The long-RNA-seq approach excludes short fragments. Consistent with our hypothesis and Fig.3A findings, 6m old APP mice were found to have substantially increased rate of B2 RNA processing compared to control mice of the same age (Fig.3E) (*p<0.05*, n=3/group).

Thus, APP mice at the active neuro-degeneration phase are characterized by higher destabilization and processing rates of B2 RNAs, consistent with the observed increase in B2 RNA regulated SRG levels in these same animals.

### Hsf1 accelerates B2 RNA processing

We then focused on the molecular mechanism underlying the increased B2 RNA processing during response to amyloid toxicity. When we had previously examined the mechanism of B2 RNA processing in non-neural cells, a member of the PRC2 protein complex, Ezh2, was reported as being responsible for the B2 RNA accelerated destabilization and processing during response to stress. However, as shown in Fig.4A-B, scant expression levels of Ezh2 levels in neural cells indicate that Ezh2 is not a key factor in B2 RNA processing in brain. Thus, the factors that may mediate destabilization for B2 RNAs during stress in neural cells remain elusive.

**Figure 4.**
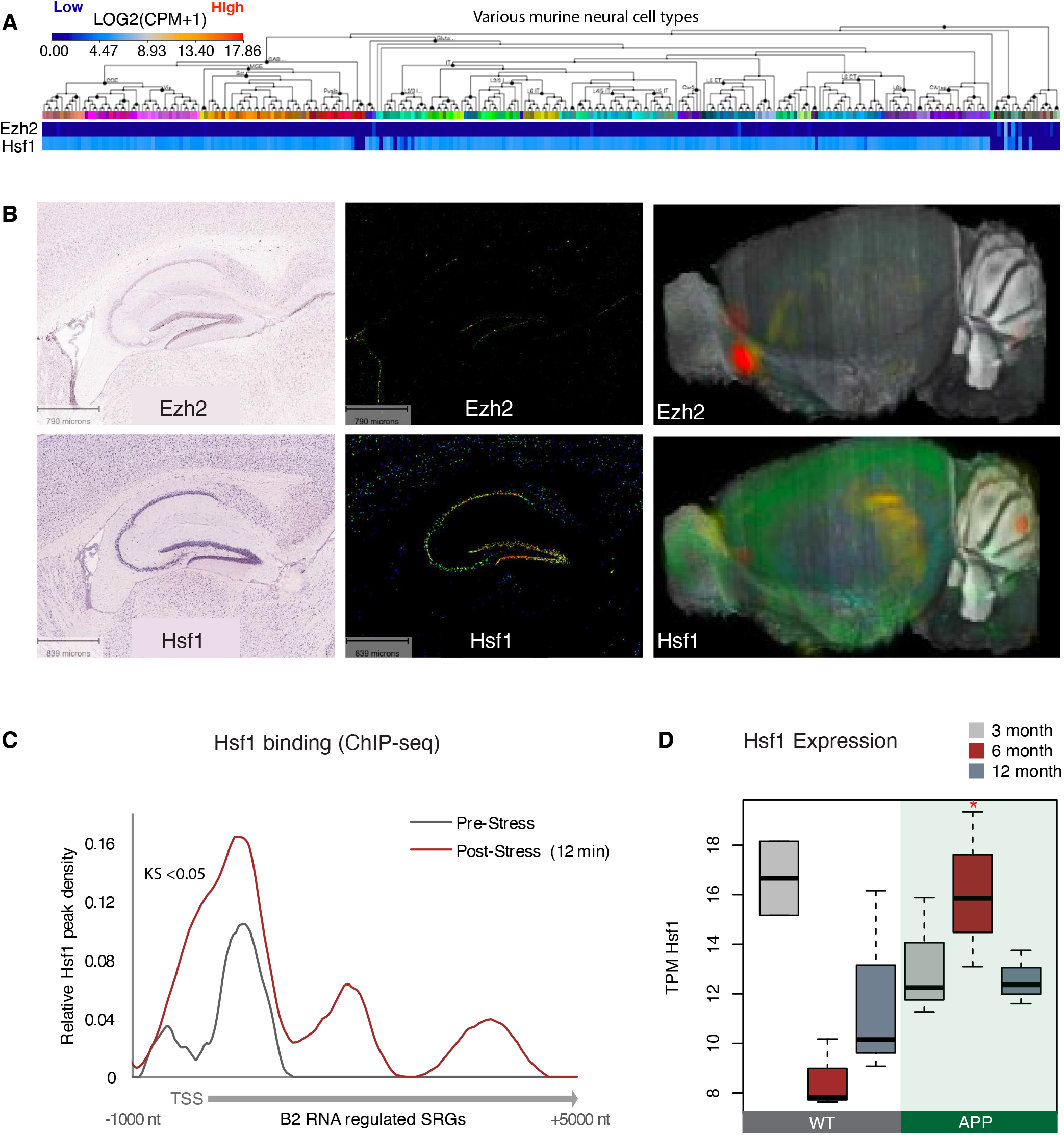
Hsf1 binds at B2 RNA regulated SRGs and is upregulated in 6m old APP mice. **(A)** Mouse cortex and hippocampus gene expression levels for Ezh2 and Hsf1 depicted in the Allen Brain Atlas Transcriptomics explorer showing limited expression of Ezh2 across multiple neural tissues compared to Hsf1. Levels are per cell type based on RNA-seq data. Image credit: Allen Institute. © [2015] Allen Institute for Brain Science. [Allen Mouse Brain Atlas]. Available from: [https://portal.brain-map.org/atlases-and-data/rnaseq#Mouse_Cortex_and_Hip]. **(B)** In situ hybridization brain image data with cellular-level resolution for Ezh2 (upper panel) and Hsf1 (lower panel) depicting low expression levels for Ezh2. Left panel, ISH images, middle panel, gene expression images, right panel, expression in 3D reconstructed whole mouse brain. Image credit: Allen Institute. © [2007] Allen Institute for Brain Science. [Allen Mouse Brain Atlas]. Available from: [https://mouse.brain-map.org/experiment/show/100142521 for Eh2 and https://mouse.brain-map.org/experiment/show/100142521 for Eh2 and https://mouse.brain-map.org/experiment/show/68196972 for Hsf1]. **(C)** Metagene analysis of increase in Hsf1 binding (ChIP-seq data) across the start of B2 RNA regulated genes (Suppl. Table 1) between pre-stress and 12 minutes post-stress (KS<0.05). ChIP-deq data are from Mahat and colleagues (Mahat et al., 2016). TSS represents the Transcription Start Site of these genes. **(D)** Boxplot depicts distribution of expression levels of Hsf1 gene among different age groups of mice between wild type and APP. Values are based on TPM counts of long-RNA-seq data. Asterisk represents p < 0.05 between 6-month old APP and 6m old WT (n=3/group, t-test).

In our earlier study that described the induction of B2 RNA processing by Ezh2, it remained unclear how Ezh2 exerted its impact on B2 RNAs, since Ezh2 lacked any known RNAse activity (Zovoilis et al., 2016). However, in subsequent experiments (Hernandez et al., 2020) we showed that instability is in fact inherent to the B2 RNA molecule and interaction with Ezh2 only accelerates this destabilization. This finding suggests that other proteins may have a similar effect on B2 RNA stability. Therefore, we started searching for stress related candidate proteins that could affect the B2 RNA processing in hippocampus.

We showed before that B2 RNA binding during response to stress is enriched near stalled RNA polymerase genomic sites. These areas are known to be highly enriched in binding sites of various stress related proteins, among which it is a stress related protein called Hsf1, which is a master regulator of stress response for various types of cellular stress (Pandey, Mandal, Jha, & Mukerji, 2011). Hsf1 has been previously connected with activation of SRGs through both transcriptional factor (TF) activities as well as other yet unknown TF-independent processes (Inouye et al., 2007). As shown in Fig.4A and 4B, in contrast to Ezh2, Hsf1 is expressed in neural tissues and especially in hippocampus. We then examined the proximity of Hsf1 binding sites, identified by the Lis lab (Mahat et al., 2016), to genes with B2 RNA binding sites (Suppl. Table 1) (Zovoilis et al., 2016). Increased Hsf1 binding was found near B2 RNA regulated SRGs (Fig.4C) and was further enriched after application of a heat-shock stimulus (KS-test < 0.05). Moreover, Hsf1 levels were found to be upregulated in APP 6m old mice compared to control group of the same age (*p<0.05*, n=3/group) (Fig.4D). As mentioned above, at the same time, 6m old APP mice have increased B2 RNA processing rates. Thus, this data suggests that Hsf1 may be a good candidate for accelerating B2 RNA processing in the context of amyloid pathology.

To check this, we investigated whether the interaction between B2 RNA and Hsf1 can accelerate B2 RNA destabilization. In particular, we incubated full length B2 RNA with the Hsf1 protein *in vitro*. As shown in Fig.5A, in the presence of Hsf1, the destabilization of full length B2 RNA was accelerated in contrast to the control protein (PNK) or no protein. No processing was observed in an RNA fragment co-incubated with B2 RNA and the protein (marked with an asterisk across Fig.5). B2 RNA destabilization in the presence of Hsf1 was accelerated compared to no protein (Fig.5B and 5C). Consistent with the above finding, the rate of processing of B2 RNA was dependent on Hsf1 concentration in the reaction (Fig.5D and 5E) and increased upon increase of Hsf1 concentration. This data shows that Hsf1 has the potential *in vitro* to accelerate B2 RNA processing.

**Figure 5.**
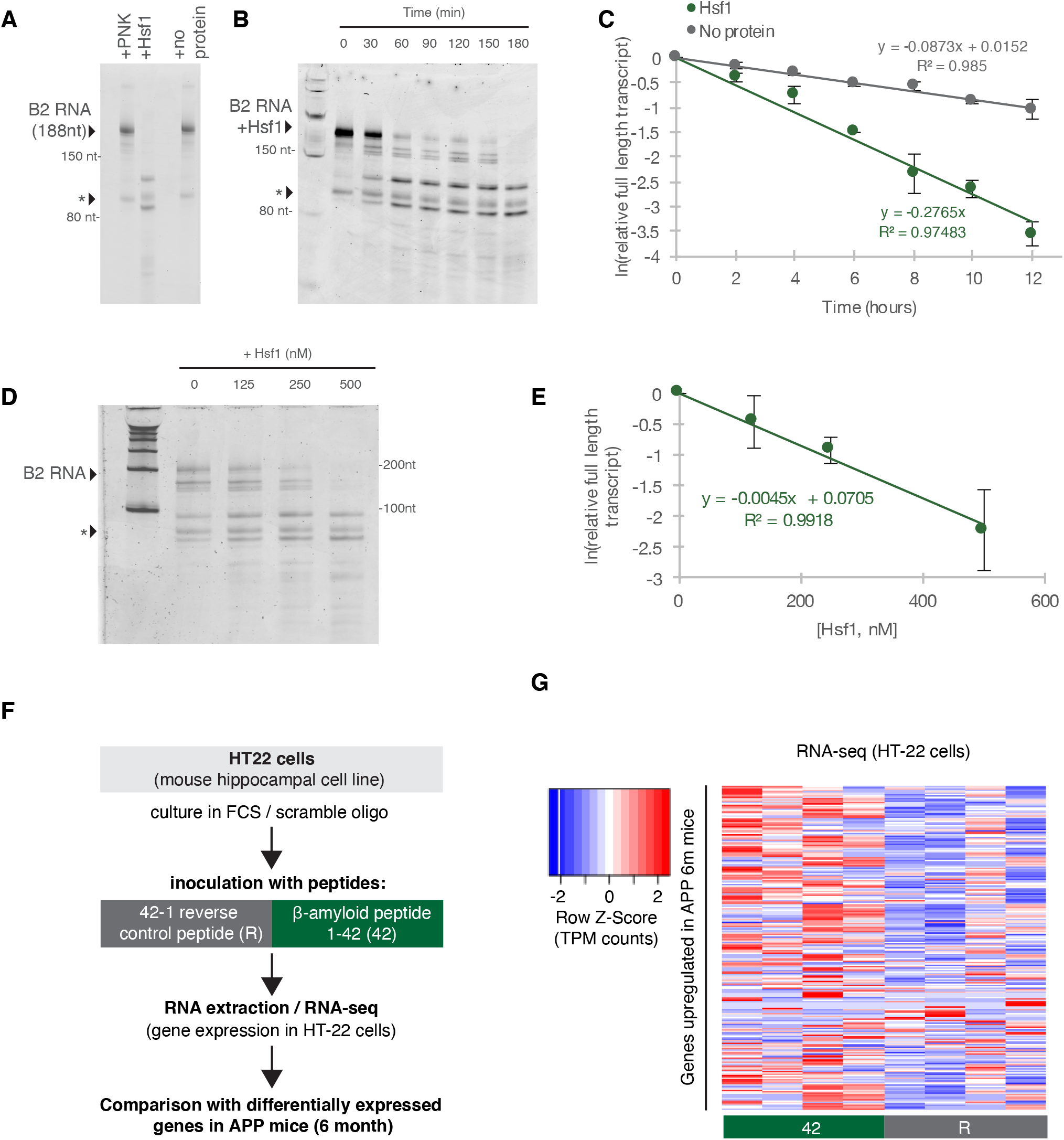
Hsf1 accelerates B2 RNA processing. **(A)** *In vitro* incubation of B2 RNA. *In vitro* transcribed and folded B2 RNA at 200nM incubated with PNK as a control (lane 1), 250nM Hsf1 (lane 2) and without protein (lane 4). Incubations occurred for 6 hours at 37°C. **(B)** *In vitro* incubation of B2 RNA for different incubation periods. *In vitro* transcribed B2 RNA (100nM) incubated at 37°C with 500nM Hsf1 in the course of 180 min with time intervals of 30 min. **(C)** Relative full length RNA remaining from (B) using ImageJ area under the curve software over time. **(D)** Titration of Hsf1 protein in incubation with 200nM *in vitro* transcribed B2 RNA. Concentrations of Hsf1 range from 0– 500nM. **(E)** Relative full length RNA remaining from (D) using ImageJ area under the curve software over time. The trendline is a linear fit, displaying standard deviation on the data points. **(F)** Experimental design for the amyloid toxicity cell culture assay employing HT-22 cells. Cells culture media were supplemented with Fetal Calf Serum (FCS) and the scramble LNA described in Figure 6 to allow comparison with the LNA experiments. **(G)** Heat map showing gene expression changes during incubation with amyloid peptides in HT22 cells. We focused on genes of Suppl. Table 2 that are genes found to be upregulated in 6m old APP mice vs 6m old WT mice to test whether our cell culture system can simulate the transcriptome observed in amyloid pathology in vivo. Heatmap depicts increase in expression values calculated by long-RNA-seq (TPM values) in the HT22 samples transfected with 1-42 (42) vs 42-1 reverse control peptide (R) (heatmap columns), for those genes up-regulated in 6m old APP mice (Suppl.Table 2) that are expressed in HT-22 cells (heatmap rows). TPM values are normalized per row. Red color represents higher expression.

### Hsf1 mediates increased B2 RNA processing in response to amyloid beta toxicity

The increased levels of Hsf1/B2 RNA processing in APP 6m old mice raised the question whether response to amyloid toxicity in hippocampal cells is connected with an Hsf1-mediated increase in B2 RNA processing. In order to test this, we employed the a hippocampal cell culture model using the a HT-22 cell line, which has been used extensively in the past as hippocampal cell stress model (Davis & Maher, 1994; Liu, Li, & Suo, 2009). We incubated these cells with amyloid beta peptides (1-42 aa) and compared their transcriptome to the one of cells incubated with an inverted sequence control peptide (R, reverse 42-1) (Fig.5F). Incubation of these cells with 1-42 amyloid beta peptides results in upregulation of a number of genes (Fig. 5G), that are also found upregulated in the 6m old APP mice (Suppl. Table 2). The increase in expression levels of genes associated with amyloid pathology suggests that this cellular model simulates to a certain extend the amyloid toxicity effect on the transcriptome of hippocampal cells.

In order to test the impact of amyloid toxicity to Hsf1-mediated B2 RNA processing we treated HT22 cells with either an LNA against Hsf1 or a scramble LNA (control) followed by incubation with the 1-42 peptides, that subject the cells to amyloid toxicity stress, or the respective control peptide (Fig. 6A). As shown in Fig.6B, treatment with the anti-Hsf1 LNA has suppressed the increase of Hsf1 levels observed in the control cells upon application of the amyloid beta peptides. As observed in the APP mice, amyloid beta peptides resulted in increased B2 RNA processing in control cells (with no Hsf1 knock down) suggesting that it is the toxicity of these peptides that induces the SINE B2 RNA transcriptome changes (Fig.6C). In contrast, under anti-Hsf1 LNA, application of the 1-42 peptides was unable to increase B2 RNA processing in the cells compared to the control cells (Fig. 6C). A similar effect was observed in B2 RNA regulated SRGs compared to other random genes (Fig.6D-6F).

**Figure 6.**
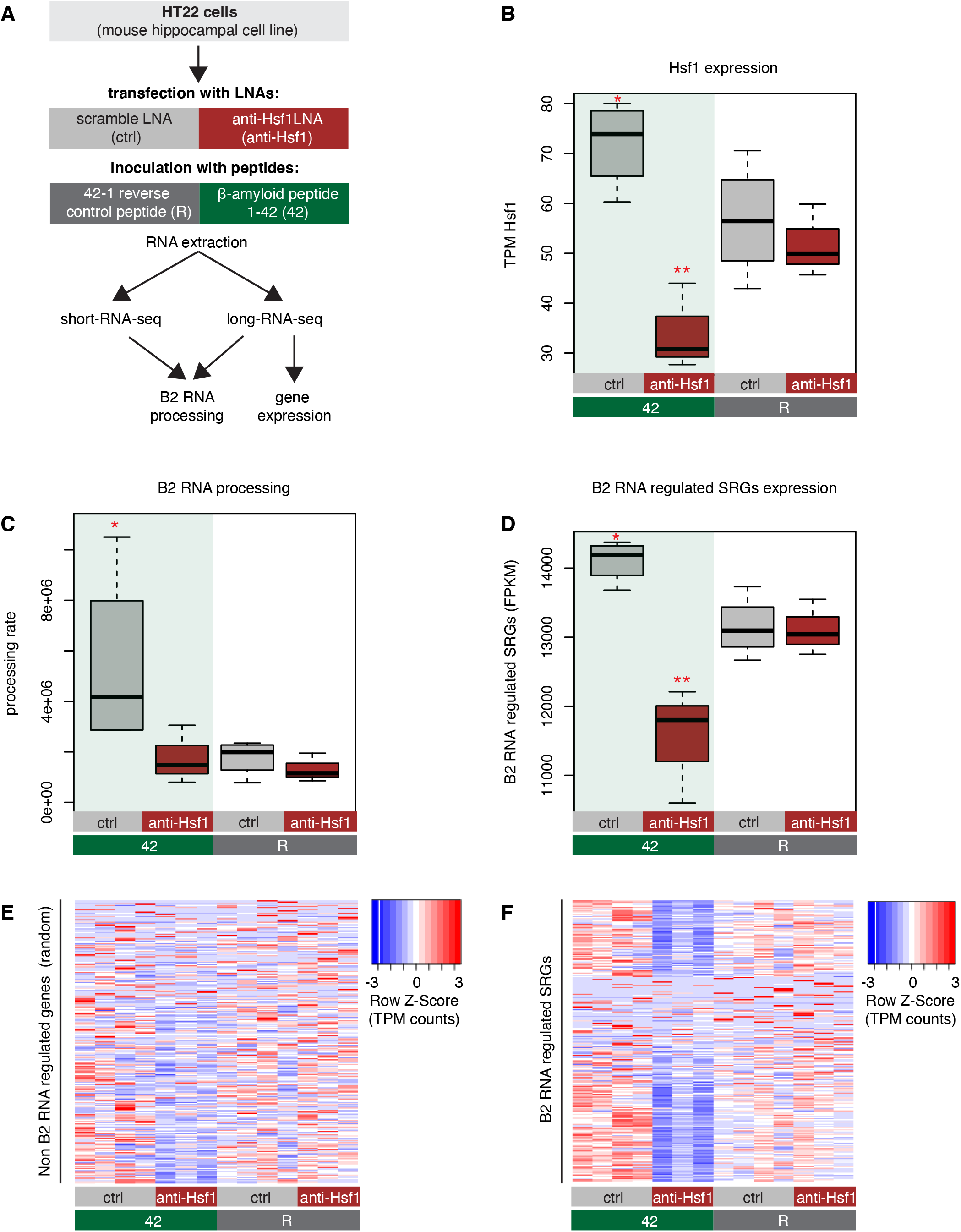
Hsf1 mediates B2 RNA processing in amyloid toxicity. **(A)** Experimental design of the combined Hsf1 Knock Down – amyloid toxicity assay in HT22 cells followed by short and long RNA-seq. **(B)** Expression levels of Hsf1 as defined by long-RNA-seq. Boxplots compare Hsf1 expression (TPM values) during incubation with the scramble LNA, and anti-Hsf1 specific LNA, incubated with either the 42 or R amyloid peptides. One asterisk represents p < 0.05 between 42/ctrl and R/ctrl (n=4/group, t-test). Two asterisks represent p < 0.05 between 42/anti-Hsf1 and all other groups (42/anti-Hsf1, n=3; R/anti-Hsf1, n=3; R/ctrl, n=4; 42/ctrl, n=4; t-test). **(C)** B2 RNA processing rate based on a combination of short and long-RNA-seq. Boxplot depicts distribution of total SINE B2 RNA processing rate among different groups of HT22 cells between 42 and R. Asterisk represents p < 0.05 between 42/ctrl and R/ctrl (n=4/group, t-test). No increase was observed between 42/anti-Hsf1 and R/anti-Hsf1 (NS, n=3/group, t-test). **(D)** B2 RNA regulated SRG expression levels based on long-RNA-seq data (FPKM values). Boxplot depicts distribution of expression levels of B2 RNA regulated genes among different groups of HT22 cells between 42 and R. Asterisk represents p < 0.05 between 42/ctrl and R/ctrl (n=4/group, t-test). Two asterisks represent p < 0.05 between 42/anti-Hsf1 and all other groups (R/anti-Hsf1, n=3; R/ctrl, n=4; 42/ctrl, n=3; t-test). **(E)** Gene expression levels (defined by long-RNA-seq) of a random set of genes not associated with B2 RNAs show weak or no association with Hsf1 treatment during response to amyloid toxicity in HT22 cells. Heatmap depicts gene expression with rows as random genes (non B2 RNA regulated) and columns corresponding to the different HT22 cell treatments. Expression values are normalized per row and correspond to TPM values, with red color represents higher expression. **(F)** Gene expression levels (long-RNA-seq) of B2 RNA regulated SRGs (Suppl. Table1) show strong association with Hsf1 treatment during response to amyloid toxicity in HT22 cells. Heatmap depicts gene expression with rows as B2 RNA regulated genes (Suppl. Table 1) and columns as different HT22 cell treatments. TPM values are normalized per row. Red color represents higher expression.

These results suggest that amyloid beta toxicity induces B2 RNA processing also *in vitro* and Hsf1 comprises a necessary component in the upstream activation pathway of B2 RNA processing and, thus, of the genes regulated by B2 RNA.

## DISCUSSION

Fewer than five ribozymes have been identified in mammals, including our previous work on RNAs made from two retrotransposon families, murine SINE B2 RNAs and human SINE ALU RNAs, which are self-cleaving RNAs (Hernandez et al., 2020). We previously showed that cleavage in SINE B2 RNAs controls response to cellular stress through activation of stress response genes. However, no connection between this novel molecular mechanism and pathologic processes was until now known. Moreover, since B2 RNAs are intrinsically reactive, and contact with Ezh2 only accelerates cleavage, it remained plausible that other stress-related proteins may also have a similar effect on accelerating B2 RNA processing, which would link this ribozyme-like property to stress response through pathways other than Ezh2. This is especially relevant in mouse tissues, such as the brain, where Ezh2 expression is limited, an expression pattern observed also in human (Suppl. Fig. 2).

Here, we unveil increased processing of SINE B2 RNAs as a novel type of transcriptome de-regulation underlying amyloid beta neuro-pathology. Our data provides a new link in the murine hippocampal pathways connecting amyloid beta toxicity with transcriptome changes in SRGs through processing of B2 RNAs. In particular, the B2 RNA processing rate increases upon progression of amyloid pathology both in mouse hippocampus and a hippocampal cell culture model, and B2 RNA regulated SRGs become hyper-activated. Consistent with the spatial proximity between B2 RNA regulated SRGs and Hsf1 binding sites, Hsf1 proved to be key for mediating B2 RNA processing in response to amyloid toxicity. Our work assigns to Hsf1 a new function that is independent of its long-established transcription factor function and includes the interaction with and processing of SINE B2 RNAs. The high levels of Hsf1 trigger a downstream cascade of events which are orchestrated into a cell-wide, SRG-mediated response to stress conditions. This axis is mediated by the ability of B2 RNA to get processed and, thus, act as a molecular switch. While healthy cells and animals are able to restore the expression levels of Hsf1, SRGs and their regulating B2 RNAs upon removal of the stress-generating stimulus, the amyloid beta load in our biological models acts as a continuous stimulus that causes the Hsf1 - B2 RNA - SRG axis to “lock” into an activated mode. Upregulation of SRGs results in increased p53 levels that induce neuronal cell death (Fig.7).

**Figure 7.**
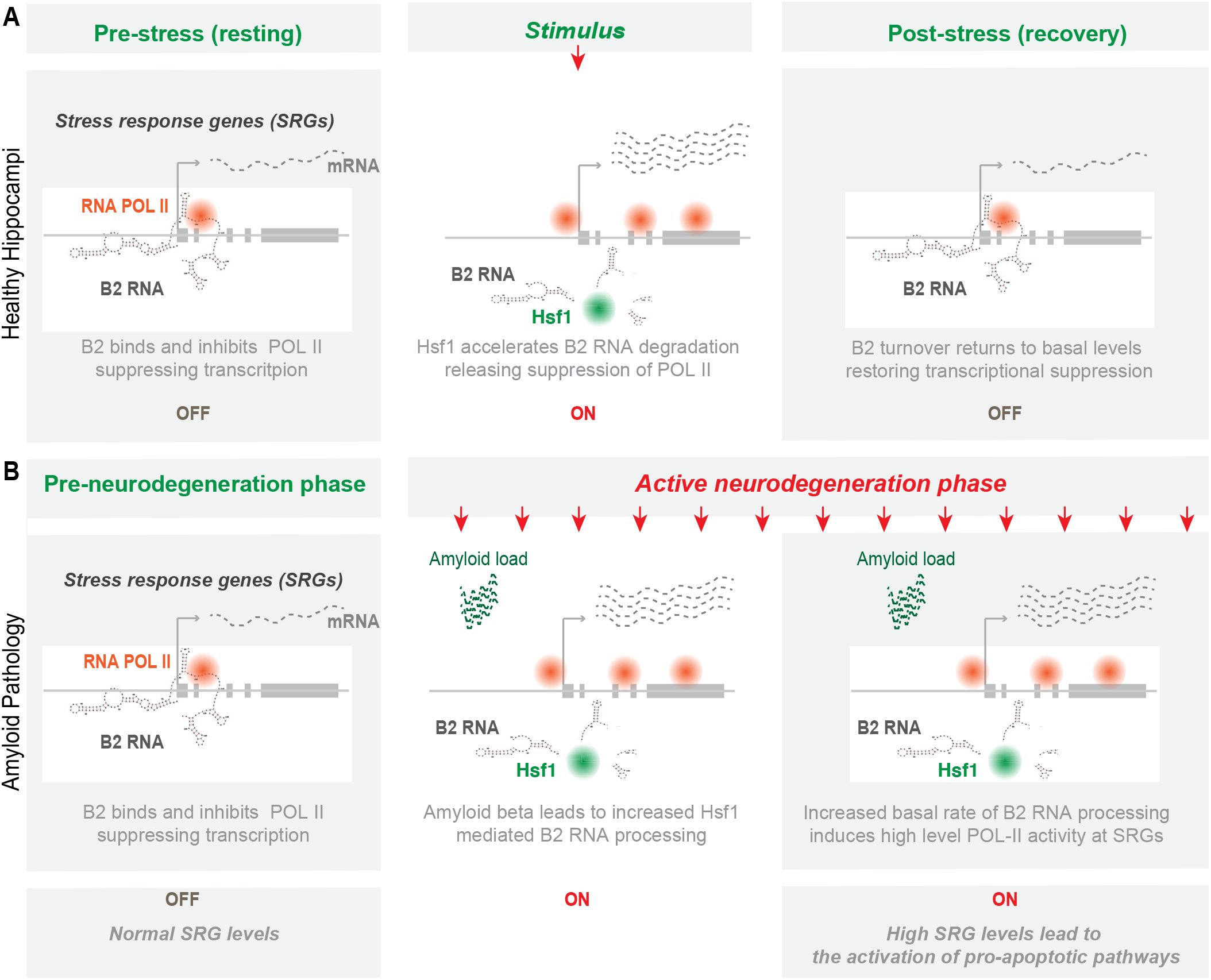
Representation of the role of B2 RNA processing in amyloid pathology. Upon removal of the stress-generating stimulus, healthy cells restore the expression levels of Hsf1, SRGs and processing rate of B2 RNAs returns to base levels. In contrast, in amyloid pathology, increased amyloid beta load acts as a continuous stimulus that causes the Hsf1 - B2 RNA - SRG axis to “lock” into an activated mode. ON/OFF represent active and suppressed SRG transcription, respectively.

In this study, we employed a mouse model of amyloid pathology in order to test the impact of increased amyloid beta load on B2 RNA processing *in vivo*. This mouse model has the NL-G-F mutations in the amyloid precursor protein knocked-in to a C57BL genomic background, with each mutation contributing to an increased severity and speed manifestation of the disease (Mehla et al., 2019). The combined effect of the APPNL−G−F mutations results in mice that experience rapid onset of AD-like symptoms at approximately 6 months (Mehla et al., 2019). This is exactly the time point that we observed the massive hyper-activation of B2 RNA SRGs, p53 and B2 RNA processing, suggesting that these changes constitute a molecular signature for the active neurodegenerative phase months, before the end state of 12 months that the mice develop terminal AD-like pathology and symptoms. Interestingly, these molecular changes were not observed in younger mice suggesting the existence of a yet unknown protective mechanism against increased neuronal activity of SRGs in 3m old APP mice. This may be attributed to the increased neuronal plasticity observed in younger brains (Lilja et al., 2013), which suggest that mechanisms in the younger brain may exist for counter-acting this excessive activity. In contrast, during aging, SRG activity in WT mice ramps down, which is not the case in APP mice, explaining at the molecular level the increased p53 levels and subsequent activation of cell death (Fig.1F).

Our study leaves a number of open questions. Particularly, our study raises the question whether a similar mode of regulation of SRGs by SINE RNAs may exist also in human and which SINE RNAs could play such a role. Similarly to B2 RNA, such SINE RNAs would be able to bind and inhibit RNA Pol II, and would be subject to a similar RNA processing mechanism enabling the release of RNA Pol II. A number of studies have described that in human, repetitive SINE RNAs of the Alu class are also upregulated during cellular stress and can bind RNA Pol II inhibiting the transcription of target genes (Yakovchuk et al., 2009). Alu RNAs are widely regarded as the equivalent in human of B2 RNA. Most importantly, as we showed before, human Alu RNAs, alike B2 RNAs, are self-cleaving RNAs and can become destabilized *in vitro* (Hernandez et al., 2020). It remains unknown whether SINE RNAs and Hsf1 play a similar role in amyloid pathology in the case of humans and whether we can extrapolate the generated conclusions in murine models to deduce that SINE RNAs are key components of the pathophysiological mechanisms underlying debilitating diseases such as AD. One major limitation compared to human pathophysiology is that the phenotype of amyloid pathology is not observed in mice even during aging. Nonetheless, a stress-central role of Alu RNAs, the human counterpart of B2 RNAs is plausible and, thus, future studies need to elucidate whether Alu RNA processing is also hyperactivated in the brain of patients with amyloid pathology in the context of AD.

Since B2 RNA regulated SRGs are highly interconnected and interweaved into various pathways, the impact of SRG hyperactivation by B2 RNA processing may extend to pathways that lie downstream of SRGs and affect various gene programs without binding directly B2 RNA. However, here it should be noted that high levels of Hsf1 are certainly expected to affect transcription also through Hsf1’s conventional transcriptional factor function while there are still many SRGs that are not directly regulated by B2 RNA. Thus, B2 RNA processing described here does not constitute the only one but just one of the parameters in the equation of SRG regulation. It remains unclear which is the interplay between Hsf1 traditional transcription factor activities and its ability to affect B2 RNA processing. Moreover, given how easily B2 RNA is processed in the presence of certain proteins, Hsf1 may be only one of the factors accelerating B2 processing in mouse hippocampus as we are just beginning to understand the implications of this form of SINE RNA regulation in cells. A broader role of SINE RNA processing in brain physiology and pathophysiology constitutes, thus, a significant possibility that could further revise our understanding of these RNAs as something more than just transcriptional noise and “junk DNA” products.

## MATERIALS AND METHODS

### Amyloid beta peptide preparations

The amyloid beta 1-42 peptides and the respective control peptides (having the reverse aa sequence compared to 1-42 peptides) were synthesized by Sigma Aldrich’s custom synthesis service using the following sequences: DAEFRHDSGYEVHHQKLVFFAEDVGSNKGAIIGLMVGGVVIA (1-42) and AIVVGGVMLGIIAGKNSGVDEAFFVLKQHHVEYGSDHRFEAD (Reverse). Upon receiving, the peptides were dissolved in 10% NH4OH at a concentration of 2.1mg/mL, sonicated for five minutes, aliquoted, dried and stored at −80ºC for further use (see below).

### Animals and behavioural measurement

Immunohistochemistry and behavioral studies of mice used are described in our previous study (Mehla et al., 2019). In brief, mouse pairs of APP-KI mice carrying Arctic, Swedish, and Beyreuther/Iberian mutations (APPNL-G-F/NL-G-F) were gifted by RIKEN Center for Brain Science, Japan, and the colony of these mice was maintained at Canadian Center for Behavioral Neuroscience vivarium. C57BL/6J mice were used as a WT control and all animals were housed in groups of 4 mice in each cage in a controlled environment (22 °C–25 °C, 50% humidity and a 12-hour light:dark cycle). All experimental procedures were approved by the institutional animal care committee and performed in accordance with the standards set out by the Canadian Council for Animal Care. We used 10 mice for behavioral tests and 2-4 mice for immunohistochemistry at each of the testing age points. Mice were individually handled each day for 1 week before starting the acquisition training and were given 8 days of acquisition trials. On each acquisition day, mice received 4 training trials from each quadrant randomly distributed each day. The latency to reach the platform was analyzed to confirm the learning behavior of the mice. A single probe trial was conducted on day 9 to assess the integrity and strength of spatial memory 24 hours after the completion of the last trial of the acquisition phase. We analyzed the data in the probe trial by measuring the time spent by mice in the target quadrant and average proximity to the escape annulus.

### Cell culture and transfections

HT22 cells, from an immortalized mouse hippocampal cell line (Davis & Maher, 1994) (Sigma Aldrich, SCC129), were cultured in DMEM (Sigma) and 1% Penicillin/Streptomycin (Gibco), Cells were thawed and passaged at least three days before the transfection date in order to allow sufficient time for cells to recover from the stress of cryopreservation and not interfere with the assessment of cellular response to stress in subsequent experiments. For knocking down of Hsf1 mRNA levels, we used an LNA long RNA GapmeR against Hsf1 (Exiqon/Qiagen) with the following sequence: 5’-CGAAGGATGGAGTCAA-3’ and an LNA long RNA GapmeR with the following scramble sequence: 5’-CCTCAATTTTATCAC-3’. The day of LNA transfections, following 5min incubation with TrypLE™ Express Enzyme (Gibco) (1x), cells were passaged and transferred to a 6-well plate at a 100,000 cells/well density and LNA transfections were performed simultaneously, using the HiPerfect reagent (Qiagen). Transfection was performed as follows: Firstly, LNAs were reconstituted in nuclease free water at a 50μM. Subsequently, 3μL 50uM LNA were mixed and incubated with 4uL Hiperfect reagent and 30ul nuclease free water, at room temperature for 20 min, and then added drop-wise to cells that had just been plated in 1ml of medium/well (still not attached) to a final LNA concentration of 150nM. Subsequently, incubation with amyloid beta (1-42) and control peptides (Reverse) was performed 24h after transfection with LNAs. Peptides were initially dissolved in DMSO and then added to cells to a final concentration of 30uM for incubation at 37C for 1 hour before treating with 0.5mL TrypLE™ (Gibco), pelleting cells at 1000rpm for 5min and resuspending the pellet in 1mL Trizol reagent (Thermofisher) for RNA extraction based on Manufacturer’s instructions.

### RNA *in vitro* transcription and RNA-protein incubations

B2 template for *in vitro* RNA transcription was ordered as IDT g-block™: 5′-GGGGCTGGTGAGATGGCTCAGTGGGTAAGAGCACCCGACTGCTCTTCCGAAGGTCCGGAGTTCAAATCCCAGCAACCACATGGTGGCTCACAACCATCCGTAACGAGATCTGACTCCCTC TTCTGGAGTGTCTGAAGACAGCTACAGTGTACTTACATATAATAAATAAATAAATCTTTAAAAAAAAA-3′. The template was amplified by PCR using a T7 promoter sequence as the forward primer: 5′-TAATACGACTCACTATAG and the following sequence as reverse primer: 5′-TTTTTTTTTAAAGATTTATTTATTTATTATATGTAAGTACA. Primers were diluted to 20mM and PCR was performed using the NEB Q5 polymerase, Q5 reaction buffer (10x), Q5 high GC enhancer (10x). The reaction proceeded at hot start 98C – 30s, (98C – 5s, 58C – 10s, 72C – 10s) X35 cycles, 72C – 10min. The samples were then analyzed by agarose gel electrophoresis (Bio-Rad, 1613100EDU) and the bands were gel extracted at the desired size and purified using the BioBasic EZ-10 gel extraction kit (BS353). A subsequent PCR was then repeated and 1ug of the amplified g-block was then *in vitro* transcribed by T7 RNA polymerase (NEB, M0251) for 2 hours at 37°C. The reaction was buffered using the T7 RNA polymerase buffer in addition to 10mM NTPs (ATP: P1132, CTP: P1142, GTP: P1152, UTP: P1162) in a final 20uL reaction. RNA was purified using the Zymo Research RNA Clean and Concentrator - 25 kit.

B2 RNA re-folding and B2 RNA – Hsf1 incubations were performed as described previously (Zovoilis et al., 2016). In brief, *in-vitro*-transcribed B2 RNA was folded with 300mM NaCl through incubation for 1 min at 50°C and cooling at a rate of 1°C/10sec until 4°C. Subsequently, B2 RNAs were incubated at a final concentration of 0.2 µM unless otherwise stated with the addition of Hsf1 diluted in TAP buffer (final reaction concentrations: 5nM Tris pH 7.9, 0.5mM MgCl2, 0.02mM EDTA, 0.01% NP40, 1% glycerol, 0.2mM DTT). Hsf1 protein incubations were performed with phosphorylated, recombinant, His-tagged Hsf1 (Enzo Life Sciences: ADI-SPP-902-F). Hsf1 working concentrations were 250nM unless otherwise specified, diluted in TAP buffer.

Fragmentation of B2 RNA was analyzed on 8M urea 10% PAGE gels stained by SYBR II (Invitrogen, S7564). Gel analysis occurred on Amersham Typhoon instruments. Band absorbance was analyzed using ImageJ area under the curve software and normalized by the ratio of experimental over initial as described previously (Zovoilis et al., 2016).

### Short-RNA-seq and long-RNA-seq

1.5μg total RNA was size separated into short and long fractions using the miRvana™ miRNA size selection kit (Thermo Fisher) as described before (Zovoilis et al., 2016). In brief, following addition of the lysis/binding buffer and the homogenate additive solution to the RNA, 1/3 of the volume 100% EtOH was added and the mix was passed through the column for binding long RNAs. 100% EtOH at 2/3 of the flow through volume was subsequently added to the flow through and passed through a second column for binding short RNAs. Eluted RNAs were tested for size and quality using the Agilent Bioanalyzer RNA pico-kit. For long-RNA-seq, the long RNA fractions were cleaned and concentrated using the RNeasy Minelute kit (Qiagen) and ribodepleted using the rRNA depletion kit (NEB). The library was then prepared using the NEBNext Ultra II direction Library preparation kit (NEB, E7760), and sample cleanups were performed using the Omega NGS Total Pure Mag Beads (Omega, SKU: M1378-01) 0.5X and 1.2X before library amplification and 0.9X following amplification. 9 cycles were used during amplification. For short-RNA-seq, the short RNA fractions were subjected to 3’-phosphoryl removal for 1 hr at 37C, treated with the T4 PNK enzyme (NEB, M0201), using exclusively the 10X PNK buffer (NEB). The short fractions were cleaned and concentrated using the RNeasy Minelute kit and the library was prepared using the NEBNext small RNA library prep set (E7330) as described before (Zovoilis et al., 2016). Sample cleanups were performed using again the Omega NGS Total Pure Mag Beads (Omega) 1.2X following library amplification. Library amplification used 15 cycles. Quantification of libraries was done by qPCR using the NEBNext library quant kit for Illumina (NEB, E7630) and library sizes were analyzed using the Agilent bioanalyzer 2100 HS DNA kit. Equimolar amounts were prepared for sequencing. Libraries were sequenced on an Illumina HiSeq platform using 150nt read lengths.

### Bioinformatics analysis

For the tissue enrichment and GO term analysis of B2 RNA regulated SRGs (Suppl. Table 1), we used the DAVID function annotation platform (DAVID 6.8, Febr. 2020) with default parameters, an EASE score of 1000 and p-adjusted values (Benjamini) (Huang da, Sherman, & Lempicki, 2009a, 2009b). For both GO Biological process and Cellular Compartment we selected the BP- or CC-direct options.

For the analysis of the short-RNA-seq and long-RNA-seq data, initially FastQC (Babraham Bioinformatics, https://www.bioinformatics.babraham.ac.uk/projects/fastqc/) was run for quality control of generated reads in fastq format. Subsequently, standard Illumina adaptor sequences were trimmed off using cutadapt-1.18 (https://doi.org/10.14806/ej.17.1.200). Short-RNA-seq reads were mapped to mouse reference genome (UCSC mm10) (November 2017) using bwa-0.7.17 in single end mode with default aln parameters (Li & Durbin, 2009). Long-RNA-seq reads for each sample were mapped to reference genome ensembl GRCm38 (November 2018) primary assembly using hisat2-2.1.0, in single end mode, with the following parameters: Report > alignments tailored for transcript assemblers including StringTie, searches > for at most 1 distinct, primary alignments for each read (Kim, Paggi, Park, Bennett, & Salzberg, 2019). SAM format files generated from mapping were converted to BAM format files using samtools-1.6 (Li et al., 2009), and to files in BED format with bamToBed utility from BEDTools-2.26.0 (Quinlan & Hall, 2010).

Models of distribution of 5’ end read fragments within the B2 loci (B2_Mm1a, B2_Mm1t and B2_Mm2) were performed using an in house python script. In brief, the script constructs a read accumulation metagene model around a hypothetical set of genomic points, in our case the start site for all B2 elements (TSS), in which the numbers of reads (or read 5' ends) around each different TSS were calculated and attributed to defined points in the model. B2 element coordinates are based on the UCSC genome browser RepeatMasker track (as of November 2018). To calculate B2 processing rate, Babraham NGS analysis suite Seqmonk 1.38.2 (https://www.bioinformatics.babraham.ac.uk/projects/seqmonk/) was used to obtain number of long reads overlapping with B2 loci (B2_Mm1a, B2_Mm1t and B2_Mm2), as well as the number of reads overlapping with tRNA loci from −5 to 15 bp. Processing rate for each sample was calculated by processed B2 count obtained from the in house python scripts normalized by tRNA from −5 to 15 bp and small reads fastq read count, as well as B2 count and long reads fastq read count: [Small fragments (position 95-110)/ [tRNAs / small RNA fastq]]/[B2RNA/long RNA fastq]

In long-RNA-seq, FPKM (Fragments Per Kilobase of transcript per Million) and TPM (Transcripts Per Million) for genes were generated using StringTie-1.3.4d (Pertea et al., 2015) with the following annotation: ensembl GRCm38 patch 94 gff3 file, and parameters limiting the processing of read alignments to only estimate and output the assembled transcripts matching the reference transcripts given in annotation and excluding non regular chromosomes. For data visualization, statistics and differential expression analysis we employed R (version 3.4.3) (https://www.R-project.org/) and the package DESeq2 (Love, Huber, & Anders, 2014). Differential expression analysis was implemented on gene count data to perform differential gene expression analysis for 6 m old mice between APP and wild type. Boxplots central line represents median and t-test was applied on the group numbers mentioned in the text.

For Hsf1 metagene analysis, we used peak.txt files of Hsf1 peaks for ChIP-seq from (Mahat et al., 2016). Peaks were analyzed with Seqmonk around Transcription start sites of genes (TSS) based on the Eponine annotation (Down & Hubbard, 2002), and filtered based on their overlap with B2 RNA regulated genes (Zovoilis et al., 2016). Also, B2 RNA binding (CHART-seq) peaks were analyzed with Seqmonk around TSS (Eponine), then filtered by overlapping with learning associated SRGs or all genes (Peleg et al., 2010). Relative density metagene plots of the distribution of the above peaks were generated using Seqmonk.

### Data access

Short and long-RNA-seq raw data have been deposited to GEO with access number GSE149243.

## ACKNOWLEDGEMENTS

This work has been supported by an Explorations Grant # 201700011 to AZ and MM from Alberta Innovates and the Alberta Prion Research Institute, a Grant # 201900003 to AZ and MM from the Alzheimer Society of Alberta and Northwest Territories and the Alberta Prion Research Institute, a Discovery Grant # RGPIN-2018-05955 to AZ from NSERC and a Compute Canada Resource Allocation Grant to AZ. AZ is supported by the Canada Research Chairs Program and the Canada Foundation for Innovation and is a former EMBO and DFG long-term fellow. YC is supported by an Alberta Innovates (AITF) fellowship. We are grateful to Dr. Angeliki Pantazi for extensively reviewing, editing and commenting on the manuscript.

## AUTHOR CONTRIBUTIONS

YC: Bioinformatics analysis, statistical analysis, establishment and testing of the analysis pipelines, data visualization and writing of the manuscript; BG: establishment and testing of the next-generation-sequencing pipelines, library construction; LS: testing of the next-generation-sequencing and in vitro assay pipelines, library construction, in vitro assays, data visualization and writing of the manuscript; CI: establishment and testing of the in vitro assay pipelines, in vitro assays; JM: acquisitions of brain tissue, hippocampal extractions; MM: design, data interpretation, writing of the manuscript; AZ: conception and design, establishment and testing of data generation and analysis pipelines, data interpretation, data visualization and writing of the manuscript, overall supervision

## CONFLICT OF INTEREST

The authors declare no conflict of interest.

---

**Supplementary Figure 1.**
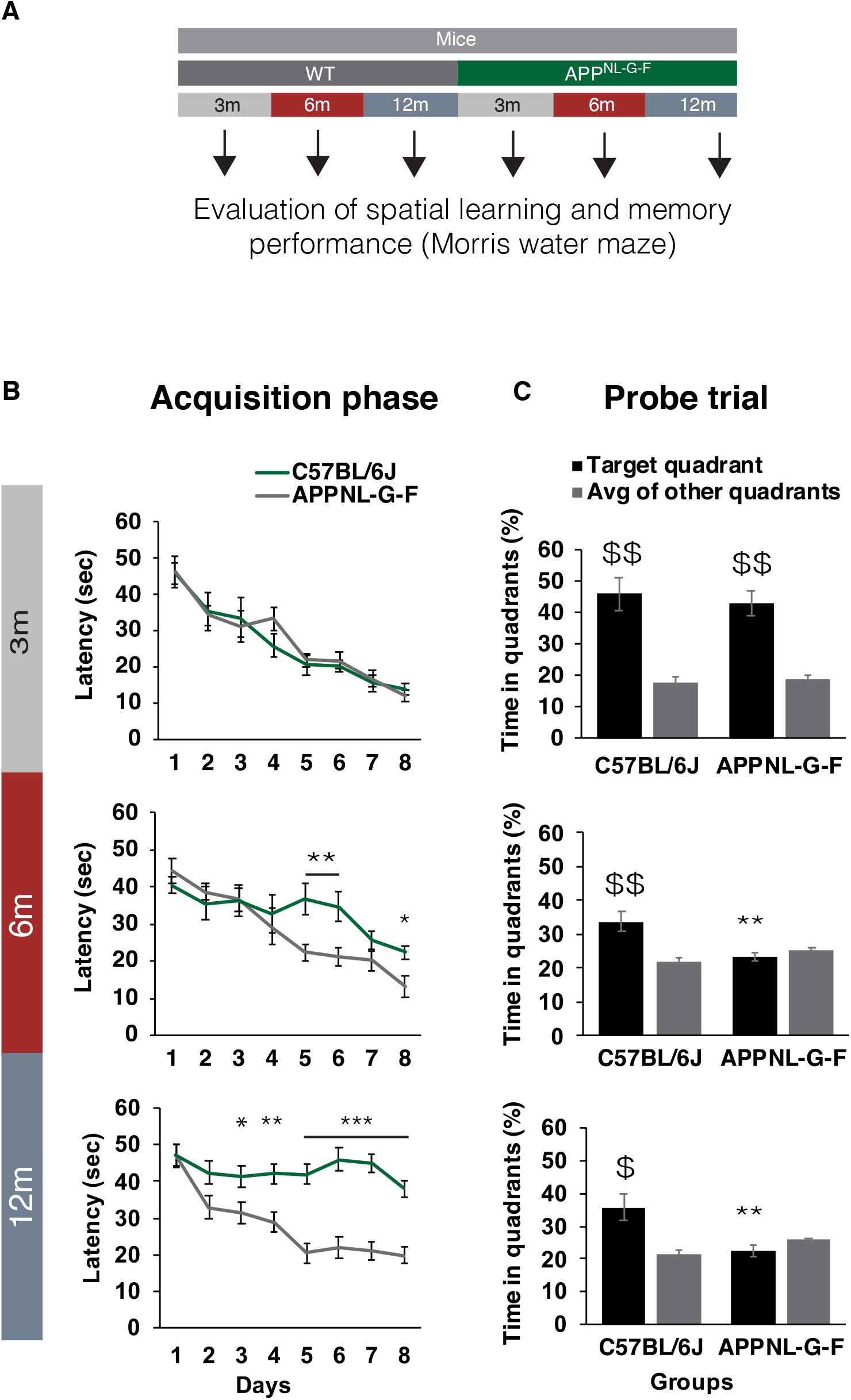
Behavioral studies (published before-adapted from Mehla and colleagues (Mehla et al., 2019)) **(A)** Experimental design. Evaluation of spatial learning and memory performance of APPNL-G-F mice used in this study in the Morris water maze based on data published in our previous study (Image credit: Mehla and colleagues (Mehla et al., 2019)). **(B)** Latency in the acquisition phase assessing the learning power. **(C)** Percent time spent in target quadrant and average of other quadrants during the probe trial. Results expressed as mean±SEM. *P < 0.05, **P < 0.01, ***P < 0.001 for the comparison between WT and APP mice; $P < 0.05, $$P < 0.01 for comparison between target quadrant and average of other quadrants (n= 10 for C57BL/6J mice; n= 10 for APPNL-G-F mice).

**Supplementary Figure 2.**
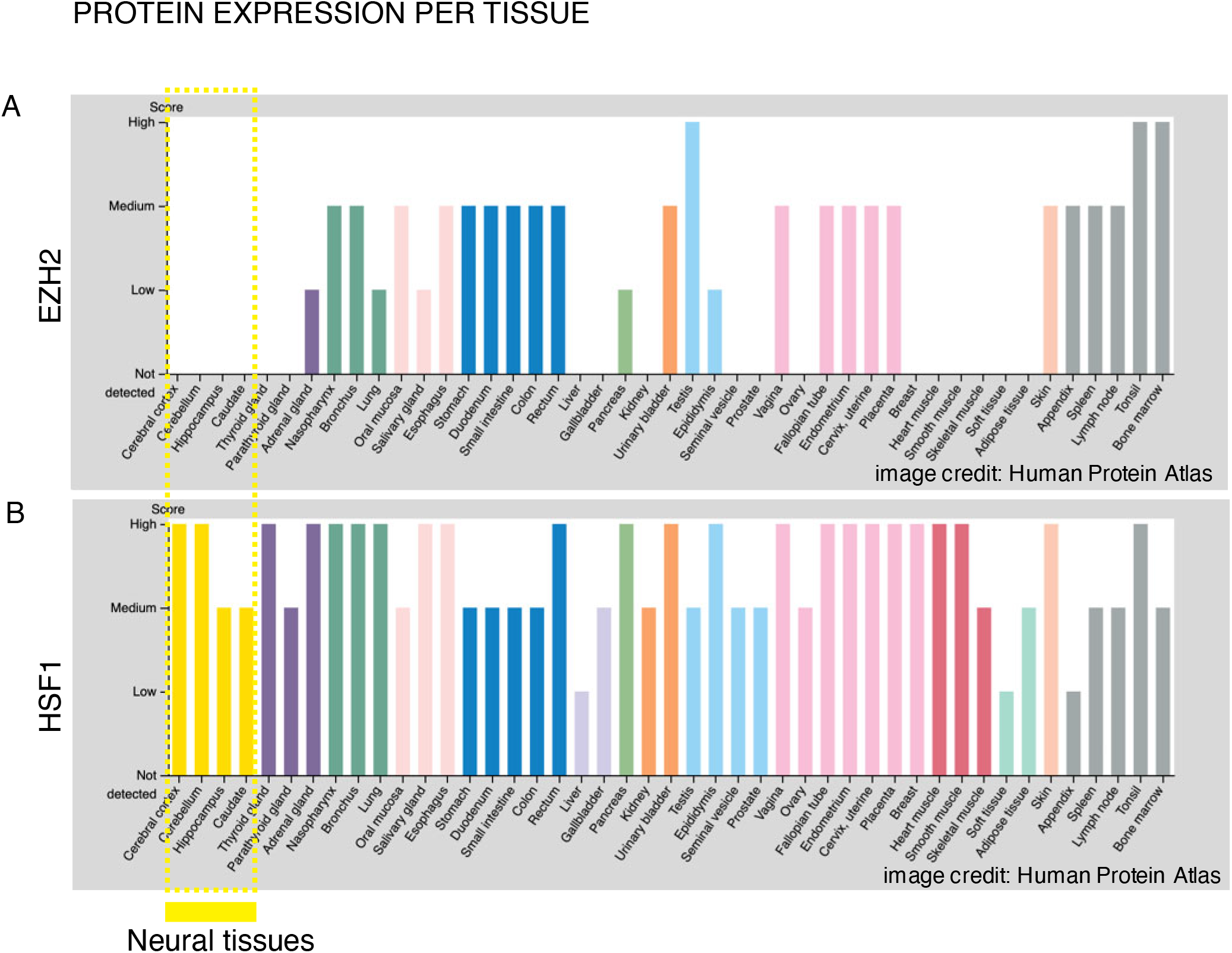
Protein expression levels of Hsf1 and EZH2 in human. Protein expression levels per tissue for Ezh2 (A) and Hsf1 (B). No expression for Ezh2 in human neural tissues is observed. Image credit: Human Protein Atlas Uhlén M et al, 2015. Tissue-based map of the human proteome. Science: PubMed: 25613900. Available from: [https://www.proteinatlas.org/ENSG00000106462-EZH2/tissue for Ezh2 and https://www.proteinatlas.org/ENSG00000185122-Hsf1/tissue for Hsf1]

